# NEO-AAV: an engineered extracellular vesicle-enveloped AAV platform for activation-coupled T cell transduction and CAR-T cell generation

**DOI:** 10.64898/2026.05.27.727849

**Authors:** Songpu Xie, Qiangbing Yang, Lisa V. Jansen, Bing Yao, Geng Yang, Kaiyong Qu, Pieter Vader, Christian Snijders Blok, Saskia C.A. de Jager, Gerard J.J. Boink, Raymond M. Schiffelers, Pieter A. Doevendans, Junjie Xiao, Joost P.G. Sluijter, Zhiyong Lei

**Author notes:** Co-Lead Authors (Songpu Xie, Qiangbing Yang).

## Abstract

Efficient delivery of genetic cargo to primary human T cells remains a critical barrier for cell-based immunotherapies. AAVs are widely used but limited by immune neutralization and poor T cell transduction efficiency. We present NEO-AAV, an engineered fusion protein (PH–ALG2–PKD12) that recruits AAV capsids into endogenous extracellular vesicles (EVs) via PI(4,5)P2-directed membrane targeting, multivalent PKD12–capsid clustering, and ALG2-mediated ESCRT machinery recruitment, yielding EV-enveloped particles with improved AAV loading compared with passive EV-AAV controls. NEO-AAV displayed enhanced resistance to an anti-AAV6 neutralizing antibody (ADK6), maintaining transduction at concentrations that neutralized naked AAV6 and outperformed passively formed EV-AAV6. Surface display of a CD7/CD3/CD28 tri-chimera enabled single-step activation and transduction of CD7⁺ T cells within PBMCs, reaching ∼38% eGFP⁺ cells without exogenous pre-activation. As a proof-of-concept, NEO-AAV generated functional CAR-T cells from primary human PBMCs, exhibiting antigen-specific IFN-γ release and cytotoxicity against CD19^+^ target cells. CAR expression peaked at day 5 and declined by day 15, consistent with episomal AAV kinetics, framing NEO-AAV as an activation-coupled delivery module rather than a durable solution. Together, NEO-AAV provides a programmable EV-enveloped AAV platform with improved immune shielding and single-step T cell transduction, offering a building block for immune cell engineering with in vivo potential.

## Introduction

Gene-based therapeutics hold enormous promise for treating a wide range of diseases, including cancer, inherited disorders, and infectious diseases ^1, 2^. A major bottleneck, however, in their clinical translation is the development of delivery systems capable of efficiently transporting therapeutic cargos into specific target cells while minimizing off-target effects, thereby inducing low immunogenicity and toxicity problems ^3–5^. Intracellular delivery of large biomolecules such as proteins, nucleic acids, and viral vectors remains challenging due to multiple biological barriers, including cellular uptake ^6^, endosomal escape ^7^, and immune clearance ^8, 9^.

Adeno-associated viruses (AAVs) have emerged as a leading vector system for in vivo gene therapy owing to their ability to mediate sustained transgene expression and a favorable safety profile ^10^. A broad range of AAV serotypes has been developed and optimized for different tissues and applications, enabling efficient gene delivery in organs such as liver, muscle, and the central nervous system ^11^. Nevertheless, systemic AAV delivery remains constrained by several fundamental limitations. High vector doses are often required to achieve therapeutic efficacy, which can lead to dose-dependent toxicities ^12^. Pre-existing neutralizing antibodies can rapidly clear circulating vectors, limiting patient eligibility and preventing redosing ^8^.

In the context of primary human T cells, efficient AAV-mediated gene delivery has remained particularly challenging. Among available serotypes, AAV6 has been reported to exhibit relatively higher transduction efficiency in human T cells ^13–15^, yet even AAV6-mediated delivery typically requires high vector doses and prior T cell activation to achieve meaningful gene transfer. This intrinsic resistance reflects several cellular properties of primary human T cells, including their non-adherent nature, limited endocytic activity, and tightly regulated membrane trafficking processes ^16^, rendering resting T cells poorly permissive to viral entry. Prior activation, such as CD3/CD28 stimulation, is typically required before T cells become susceptible to transduction, a process that can alter their phenotype and functionality ^17, 18^. Collectively, these constraints restrict the broader application of AAV-based strategies in immunotherapy and personalized cell therapy.

Compounding these entry barriers, inefficient endosomal escape and limited nuclear access of delivered genomes further restrict transduction efficiency ^7^. Together, these constraints highlight a major unmet need for delivery strategies that enable efficient, targeted, and minimally perturbative genetic modification of primary T cells, ideally with activation cues integrated into the delivery step itself, for advancing cell-based immunotherapies, including CAR-T cell generation ^19, 20^.

Extracellular vesicles (EVs) are endogenous nanoscale particles secreted by virtually all cell types and play a natural role in intercellular communication by transporting proteins, RNA, and other biomolecules ^21, 22^. Their intrinsic biocompatibility, low immunogenicity, and ability to cross biological barriers make EVs an attractive platform for therapeutic delivery ^23, 24^. However, clinical translation of EV-based therapies has been hindered by limited cargo loading efficiency and poor control over EV biogenesis, resulting in insufficient delivery efficacy ^25–27^. Naturally occurring extracellular vesicle–associated AAV (EV-AAV) particles have been reported to confer partial protection from neutralizing antibodies and modest improvements in delivery efficiency. However, in these systems, vesicle association occurs largely passively and lacks precise control over cargo loading and targeting, resulting in heterogeneous vesicle populations and variable delivery outcomes ^28–30^.

Engineering active control over EV–AAV association therefore represents a promising strategy to improve both immune evasion and delivery specificity. Building on a stepwise EV-biogenesis scaffold concept developed by our group ^31^, we sought to re-engineer AAV biogenesis by coupling viral assembly with endogenous EV secretion. Here, we present NEO-AAV, an engineered AAV envelopment construct that redirects assembled AAV capsids into EV biogenesis routes. Central to this architecture is the multivalent interaction between the AAV capsid and the AAV receptor–derived PKD1–PKD2 tandem domain (PKD12) ^32–35^. Because each AAV capsid presents multiple PKD12 binding sites, PKD12 functions as a virion-engaging multivalent clustering module that is expected to drive higher-order capsid organization and promote membrane recruitment, functionally analogous to a self-assembling scaffold. Through this engineered envelopment route, NEO-AAV establishes a synthetic AAV biogenesis pathway that couples viral assembly to EV-mediated secretion in the producer cell. Leveraging this modular platform, we further incorporate surface-display targeting modules inspired by engineered virus-like particle (VLP) systems, in which multivalent receptor-targeting and immune-engaging chimeric modules enable efficient, cell-type–specific delivery and, in the context of co-stimulatory ligand display, concurrent primary human T cell activation ^19^. NEO-AAV exhibited enhanced resistance to neutralizing antibodies in HEK293FT reporter cells and, when combined with the CD7/CD3/CD28 tri-chimera, improved transduction efficiency in CD7^+^ T cells within primary human PBMCs, which are notoriously resistant to AAV-mediated gene delivery. Using this tri-chimera surface display, we further show that functional CAR-T cells can be generated from primary human peripheral blood mononuclear cells (PBMCs) through a single activation-coupled transduction step, providing an alternative to the conventional two-step bead-activation-then-transduction workflow.

Here we describe the design, characterization, and proof-of-concept application of NEO-AAV, and delineate both what this architecture demonstrates and what remains to be established.

## Results

### An engineered envelopment architecture for EV-mediated AAV secretion

To build a stepwise envelopment strategy capable of redirecting AAV capsids into the EV secretion pathway, we distilled generalizable principles governing membrane envelopment and vesicle release from cellular biology and enveloped-virus biogenesis. Across diverse biological contexts, including EV secretion and the life cycles of enveloped viruses, productive envelopment relies on a coordinated integration of membrane targeting ^39–42^, membrane deformation ^43, 44^, and membrane scission ^45–47^. Importantly, these processes are orchestrated by host cell machinery but can be reconstituted through modular protein engineering. Guided by these principles, we designed NEO-AAV as a three-module fusion construct (PH–ALG2–PKD12) intended to operate through three coordinated steps (see Fig 1.).

**Fig. 1.**
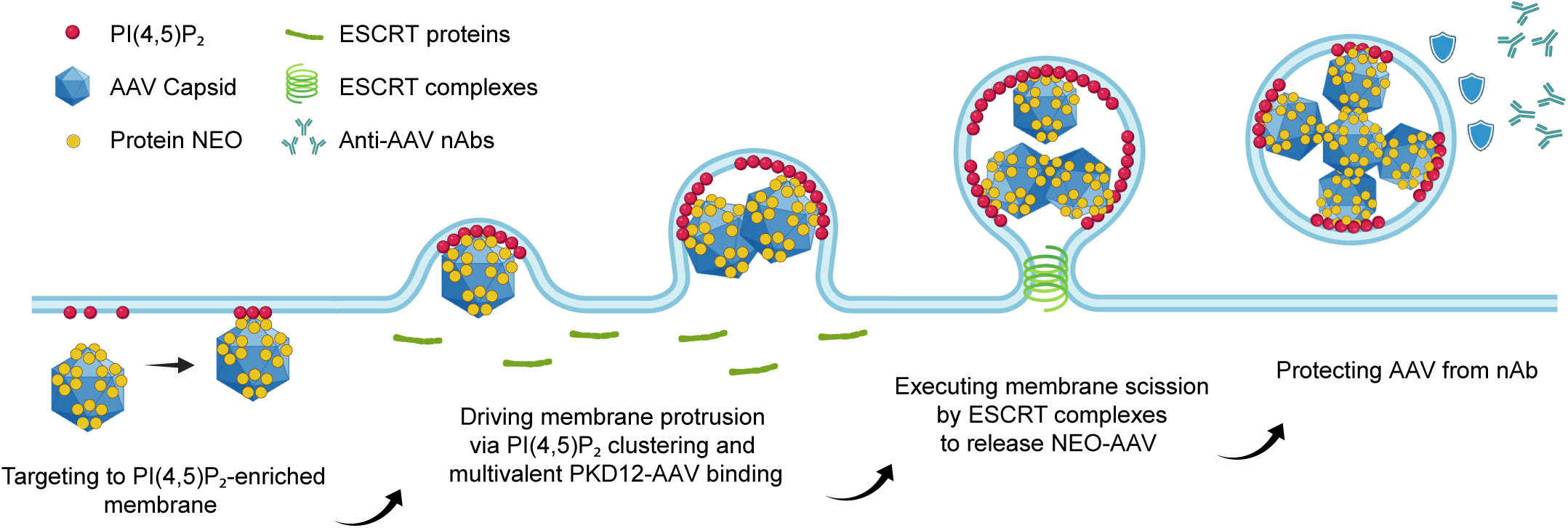
Design principles and mechanism of the NEO-AAV envelopment platform. Schematic illustration of the engineered NEO-AAV biogenesis pathway, which reprograms AAV assembly into an extracellular vesicle (EV)-mediated secretion route. The process consists of three coordinated steps: (1) membrane targeting via the pleckstrin homology (PH) domain to PI(4,5)P₂-enriched regions of the plasma membrane; (2) multivalent interactions between the PKD1–PKD2 tandem domain (PKD12) and assembled AAV capsids, generating local membrane curvature and initiating membrane deformation; and (3) ALG2-mediated ESCRT recruitment designed to facilitate membrane scission and release of EV-enveloped AAV particles. Shield icons indicate protection from neutralizing antibodies (nAbs).

#### Step 1. Membrane targeting to PI(4,5)P₂-enriched domains

Efficient vesicle formation requires the spatial concentration of relevant components at defined regions of the plasma membrane. Many EV biogenesis pathways and enveloped viruses (e.g., HIV-1 assembly mediated by Gag) initiate assembly at plasma membrane microdomains enriched in the phosphoinositide PI(4,5)P₂, which serves as a central spatial organizer for membrane-associated proteins ^39–42^. To approximate this step, we incorporated a PH domain into the NEO construct to mediate targeting to PI(4,5)P₂-enriched membrane microdomains, consistent with its well-established role as a phosphoinositide-binding module ^48^.

#### Step 2. Multivalency-driven membrane deformation

Productive membrane envelopment requires physical bending of the lipid bilayer to encapsulate large cargo assemblies. Rather than being driven solely by cytoskeletal forces, membrane deformation can arise from the local accumulation and multivalent clustering of membrane-associated proteins ^43, 44^. In viral systems, higher-order assembly of structural proteins, such as HIV Gag, or multivalent interactions between capsids and host factors can generate sufficient membrane curvature to drive budding ^39^. Leveraging this principle, the NEO construct incorporates a PKD12 domain, derived from the AAV receptor (AAVR) and capable of binding assembled AAV capsids ^32–35^, to exploit the presence of multiple PKD12-binding sites on the AAV capsid as a source of multivalency ^33, 34^, which we hypothesized would drive higher-order capsid clustering at the membrane and generate sufficient curvature to induce envelopment.

#### Step 3. ESCRT machinery-mediated membrane closure and release

The final step of envelopment requires scission of the membrane neck connecting the nascent vesicle to the plasma membrane. In cellular systems, this process is mediated by the host cell’s endosomal sorting complexes required for transport (ESCRT) machinery, which catalyzes membrane constriction and fission ^45–47^. To enable efficient vesicle release, the NEO architecture is designed to promote recruitment of endogenous ESCRT machinery at budding sites, facilitated by the ALG2 (also known as PDCD6) module incorporated in the fusion construct, a factor implicated in ESCRT machinery-mediated plasma membrane repair ^49^. Through interactions with ESCRT-associated proteins, ALG2 is expected to facilitate membrane scission, thereby converting membrane protrusions into productive vesicle release.

Collectively, this framework enables the redirection of assembled AAV capsids into endogenous EV biogenesis pathways, enabling active secretion of enveloped viral particles. These EV-enveloped AAVs are released into the extracellular space, where the surrounding membrane provides a protective barrier against anti-AAV neutralizing antibodies (Fig. 1).

### NEO-AAV produces EV-enveloped AAV via an engineered secretion route

To determine whether the engineered envelopment architecture could redirect AAV biogenesis toward extracellular vesicle secretion, we constructed a series of NEO-AAV variants incorporating functionally distinct PKD12 modules (Fig. 2A). These included the native PKD12 from the AAV receptor (AAVR; PKD12_WT_), an engineered high-affinity variant (PKD12_EN_) ^50^, and a non-binding homolog control derived from KIAA0319 (PKD12_NC_) ^35^. In contrast to conventional EV–AAV systems, where vesicle association occurs passively and with low efficiency, NEO-AAV establishes a programmed envelopment pathway through direct multivalent interactions between assembled AAV capsids and PKD12, enabling active capsid recruitment and higher-order clustering at the membrane. This panel enabled systematic interrogation of capsid-binding strength as a determinant of EV-mediated AAV envelopment, while the PKD12_NC_ variant served as a binding-deficient negative control that retains the PH and ALG2 modules.

**Fig. 2.**
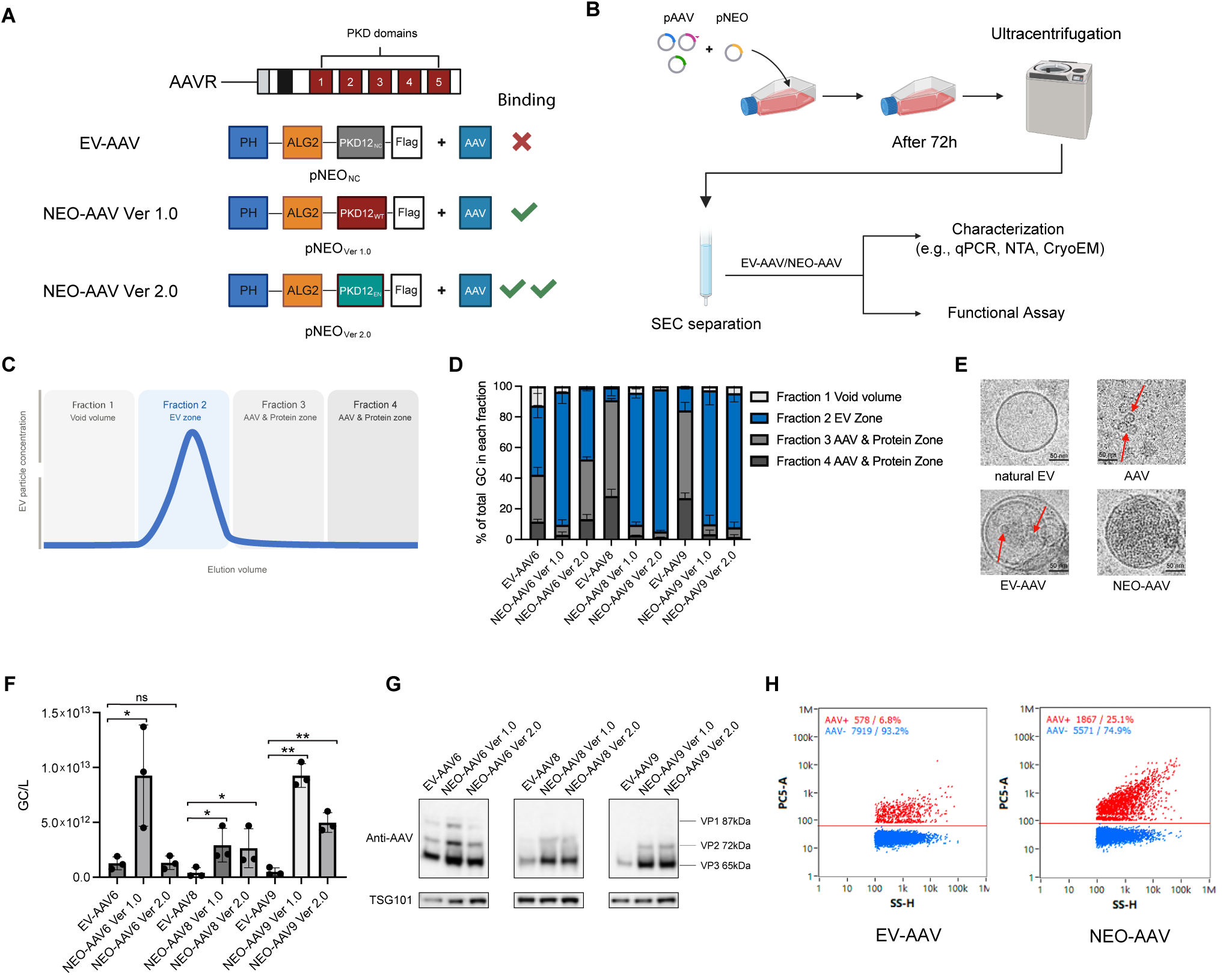
Engineering and characterization of NEO-AAV-mediated EV envelopment across AAV serotypes. (**A**) Schematic overview of EV-AAV and NEO-AAV constructs incorporating different PKD12 variants: non-binding control (PKD12NC), wild-type (PKD12WT), and engineered high-affinity variant (PKD12EN). Double checkmark indicates that PKD12EN was previously reported to exhibit higher binding affinity for AAV capsids; however, this did not translate to improved EV loading for all serotypes tested (see panel F). (**B**) Experimental workflow for production and purification of EV-AAV and NEO-AAV. HEK293FT cells were transfected with AAV packaging plasmids together with NEO constructs, followed by collection of conditioned medium at 72 h, ultracentrifugation, and size exclusion chromatography (SEC). Downstream analyses included qPCR, nanoparticle tracking analysis (NTA), cryo-electron microscopy (cryo-EM), and functional assays. (**C**) Schematic representation of SEC fractionation using a qEV column, illustrating the separation into void volume (Fraction 1), EV-enriched fraction (Fraction 2), and free AAV/protein fractions (Fractions 3–4), based on the manufacturer’s size-exclusion profile. (**D**) Distribution of AAV genome copies across SEC fractions. (**E**) Representative Cryo-EM images of natural EVs, free AAV, EV-AAV, and NEO-AAV particles. Scale bars, 50 nm. (**F**) EV-fraction viral genome yield (GC/L) across AAV6, AAV8, and AAV9 serotypes for EV-AAV and NEO-AAV variants. Data are presented as mean ± SEM from n = 3 independent biological replicates. Within each serotype, NEO-AAV variants were compared to EV-AAV controls using one-way ANOVA followed by Dunnett’s multiple comparisons test against the EV-AAV control within each serotype. ns, not significant; * p < 0.05; ** p < 0.01. (**G**) Western blot analysis of EV-associated fractions confirming the presence of AAV capsid proteins (VP1, VP2, VP3) and enrichment of the EV marker TSG101. (**H**) Nano-flow cytometry scatter plots of single EV particles from EV-AAV6 and NEO-AAV6 preparations, with AAV^+^ (red) and AAV^-^ (blue) populations indicated. AAV^+^ proportions: EV-AAV6 6.8%, NEO-AAV6 25.1%.

HEK293FT cells were transfected with standard AAV packaging components together with NEO-AAV constructs, followed by collection of conditioned medium after 72 h and purification via ultracentrifugation and subsequent size exclusion chromatography (SEC) (Fig. 2B). Following SEC purification, four major fractions were consistently observed, corresponding to void volume (Fraction 1), EV-enriched fractions (Fraction 2), free AAV, and soluble proteins (Fractions 3 and 4) (Fig. 2C).

Quantitative PCR analysis across different SEC fractions demonstrated a pronounced redistribution of viral genomes toward the EV-enriched fraction (Fraction 2) in NEO-AAV preparations compared to naturally occurring EV-AAV controls (Fig. 2D). Notably, for AAV6, NEO-AAV Ver 1.0 produced the highest absolute EV-fraction genome yield, whereas the engineered high-affinity PKD12_EN_ variant (Ver 2.0) did not improve loading — an outcome we interpret as serotype-dependent PKD engagement, to be addressed by directed evolution of capsid-binding variants in future work.

Cryo-EM analysis confirmed the vesicular identity of NEO-AAV particles and distinguished them from control preparations (Fig. 2E). Natural EVs displayed smooth membrane-bound structures with no internal capsid density, and free AAV appeared as naked capsids (∼25 nm) lacking any surrounding membrane. EV-AAV particles showed membrane association but with sparse internal capsid content. In contrast, NEO-AAV vesicles exhibited dense capsid loading within the vesicle lumen, consistent with active viral envelopment through the engineered pathway.

Consistent with this redistribution, NEO-AAV achieved substantially higher EV-fraction genome copy yields relative to EV-AAV controls across all three serotypes tested (Fig. 2F), with the most pronounced enhancement observed for AAV9, confirming genuine capsid enrichment within the vesicle fraction rather than a redistribution attributable to loss of free AAV. Immunoblot analysis confirmed robust incorporation of AAV capsid proteins (VP1–3) within EV-associated fractions, together with enrichment of canonical EV markers such as TSG101, validating the vesicular identity of NEO-AAV particles (Fig. 2G).

Nano-flow cytometric analysis of single EVs showed an increase in AAV^+^ EV populations in NEO-AAV6 samples compared to EV-AAV6 controls (Fig. 2H; 25.1% vs 6.8% AAV^+^ events), consistent with enhanced capsid packaging into secreted vesicles at the single-particle level.

Together, these results confirm that NEO-AAV establishes a programmed envelopment route that substantially enriches AAV incorporation into secreted EVs compared with passive EV-AAV association, though a large proportion of secreted vesicles remain AAV^-^, underscoring the potential for further optimization of packaging efficiency. By leveraging multivalent capsid–PKD12 interactions to drive membrane recruitment and ESCRT-dependent vesicle release, NEO-AAV creates a synthetic AAV biogenesis pathway that couples viral assembly with endogenous EV secretion.

### EV-enveloped NEO-AAV exhibits enhanced stability and resistance to neutralizing antibodies

A major limitation of systemic AAV delivery is rapid neutralization by pre-existing anti-AAV antibodies, which severely restricts transduction efficiency and prevents vector redosing. We therefore asked whether EV envelopment via the NEO-AAV pathway could functionally shield AAV capsids from antibody-mediated neutralization.

Because PKD12_EN_ did not improve AAV6 loading efficiency and behaved comparably to EV-AAV controls (Fig. 2), subsequent functional analyses were performed using NEO-AAV6 Ver 1.0 (hereafter referred to as NEO-AAV6). AAV6, EV-AAV6, and NEO-AAV6 preparations were pre-incubated for 1 hour with increasing concentrations of anti-AAV neutralizing antibodies (0, 5, 50, and 500 ng mL⁻¹) prior to transduction of HEK293FT cells at an MOI of 1.5 × 10⁴, followed by quantification of eGFP expression by flow cytometry after 72 h, as outlined in Fig. 3A.

**Fig. 3.**
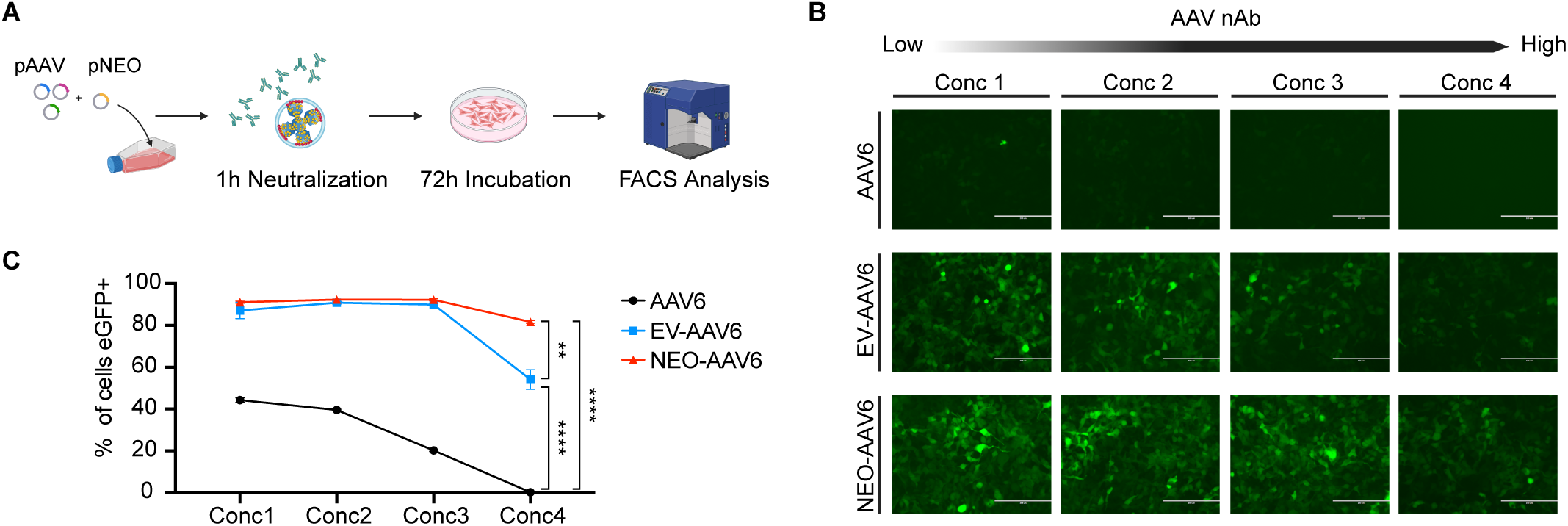
EV envelopment confers resistance to antibody-mediated neutralization. (**A**) Schematic of the neutralization assay. AAV6, EV-AAV6, or NEO-AAV6 particles were pre-incubated for 1 h with increasing concentrations of anti-AAV neutralizing antibodies (0, 5, 50, and 500 ng mL⁻¹), followed by transduction of HEK293FT cells at a multiplicity of infection (MOI) of 1.5 × 10⁴. Cells were analyzed after 72 h by flow cytometry. (**B**) Representative fluorescence microscopy images showing eGFP expression in HEK293FT cells under increasing neutralizing antibody concentrations for AAV6, EV-AAV6, and NEO-AAV6. Scale bars, 200 μm. (**C**) Quantification of transduction efficiency (% eGFP⁺ cells) across antibody concentrations for AAV6, EV-AAV6, and NEO-AAV6. Data are presented as mean ± SEM from n = 3 independent biological replicates. Statistical significance was determined by one-way ANOVA followed by Tukey’s multiple comparisons test within each antibody concentration. ** p < 0.01; **** p < 0.0001.

In the absence of neutralizing antibodies (0 ng mL⁻¹), both EV-AAV6 and NEO-AAV6 achieved approximately two-fold higher transduction efficiency compared with naked AAV6, consistent with previously reported EV-mediated enhancement of AAV delivery 28-30. With increasing antibody concentrations, naked AAV6 exhibited a progressive decline in transduction efficiency, as illustrated by representative fluorescence images (Fig. 3B). EV-AAV6 showed partial protection at lower antibody concentrations but exhibited a pronounced reduction at the highest concentration (500 ng mL⁻¹). In contrast, NEO-AAV6 maintained robust transduction across all tested concentrations (Fig. 3C), consistent with more effective EV envelopment and capsid shielding.

Together, these results demonstrate that both EV-AAV6 and NEO-AAV6 exhibit enhanced transduction efficiency compared with naked AAV6 under antibody-free conditions. Under increasing antibody pressure, NEO-AAV6 demonstrated superior resistance compared with EV-AAV6, with the difference becoming most pronounced at the highest antibody concentration tested (500 ng mL⁻¹). These findings suggest that active EV envelopment via the NEO-AAV pathway confers additional shielding capacity beyond that of passively formed EV-AAV, though validation against polyclonal human sera will be required to establish the generalizability of this effect.

### Targeted NEO-AAV delivery enables efficient genetic modification of CD7^+^ T cells in primary human PBMCs

Efficient genetic modification of primary human T cells remains a major bottleneck for cell-based immunotherapies due to the intrinsic resistance of resting T cells to viral transduction. We therefore selected AAV6 as a clinically relevant benchmark serotype, given its reported permissiveness for primary human T cell transduction, albeit at typically low efficiency and requiring prior activation ^13–15^.

To enable targeted delivery and simultaneous immune activation, we engineered a modular T cell–targeting tri-chimera comprising CD7-targeting, CD3-engaging, and CD28 co-stimulatory modules ^51^, inspired by previously reported enveloped delivery strategies for in vivo T cell engineering ^19^ (Fig. 4A). The CD28-binding domain was derived from this system, whereas the CD3 scFv (OKT3) and CD7 nanobody were selected in-house. This architecture enables CD7-mediated targeting, together with CD3-mediated activation and CD28 co-stimulatory signaling, thereby coupling viral delivery with functional T cell stimulation.

**Fig. 4.**
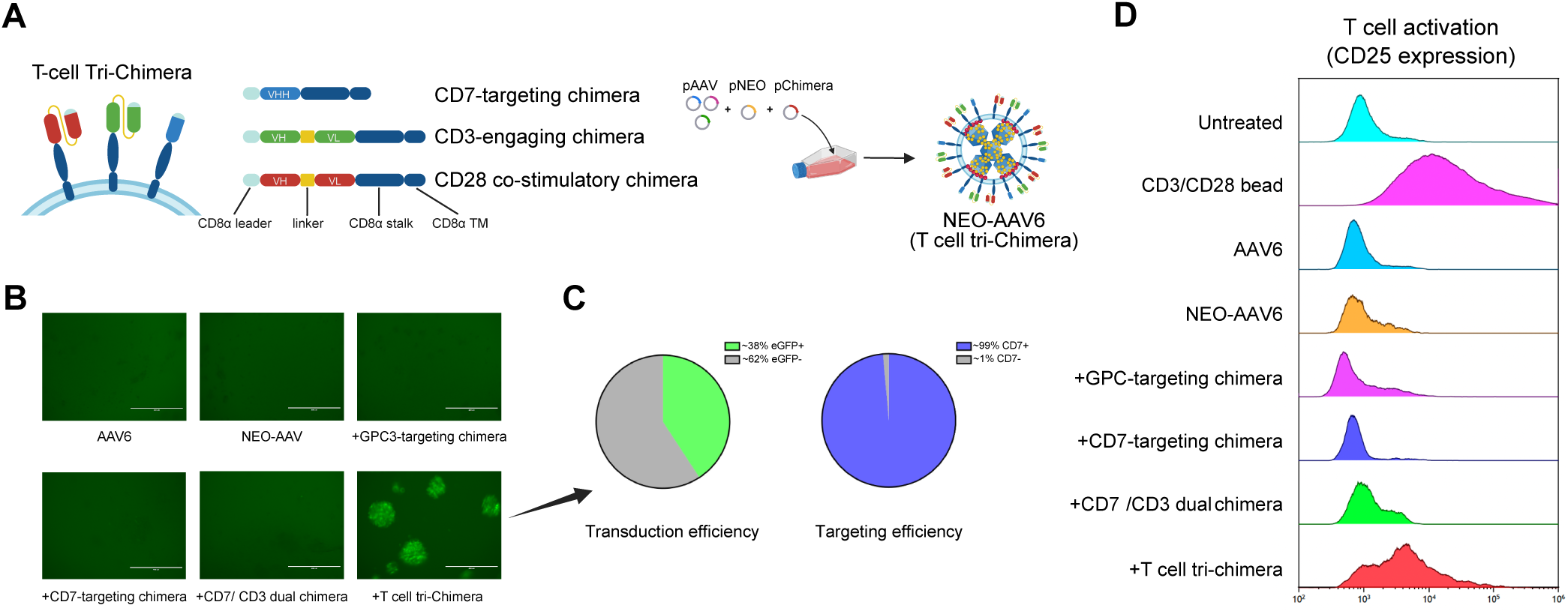
Programmable surface display of targeting and co-stimulatory chimeras enables single-step genetic modification and activation of CD7^+^ T cells in primary human PBMCs. (**A**) Schematic representation of engineered targeting chimeras incorporated into NEO-AAV, including a CD7-specific nanobody (VHH), a CD3-specific scFv, and a CD28 co-stimulatory domain. These modules are combined into a tri-chimera design for simultaneous CD7-mediated targeting, CD3-mediated engagement, and CD28 co-stimulation. (**B**) Representative fluorescence microscopy images of primary human PBMCs transduced with AAV6, untargeted NEO-AAV6, or NEO-AAV6 displaying individual or combinatorial targeting chimeras, as indicated. Scale bars, 200 μm. (**C**) Quantification of transduction efficiency (% eGFP⁺ cells) and CD7 targeting specificity (% CD7⁺ among eGFP⁺ cells) in the tri-chimera condition. Data represent mean ± SEM from n = 3 independent donors. (**D**) Flow cytometry histograms showing CD25 surface expression in total PBMCs across treatment conditions, including untreated, CD3/CD28 bead stimulation, AAV6, untargeted NEO-AAV6, and NEO-AAV6 displaying individual, dual, or tri-chimera modules. For all conditions, viral preparations were pre-incubated with anti-AAV neutralizing antibodies (500 ng mL⁻¹) prior to transduction at a multiplicity of infection (MOI) of 1 × 10⁶.

Human PBMCs were transduced with eGFP-expressing AAV6, untargeted NEO-AAV6, or NEO-AAV6 displaying individual or combinatorial chimeric modules, including a GPC3-targeting chimera as a non–T cell–targeting specificity control, followed by assessment of eGFP expression. Conventional AAV6, untargeted NEO-AAV6 and NEO-AAV6 displaying single or dual modules (CD7 or CD7/CD3) exhibited minimal transduction within the CD7^+^ T cell population.

Only presentation of the full T cell tri-chimera (CD7/CD3/CD28 modules) resulted in robust eGFP expression, accompanied by the formation of prominent eGFP^+^ T cell clusters (Fig. 4B). Under this condition, on average ∼38% of CD7^+^ cells were eGFP^+^ across independent experiments. As negligible transduction was observed in all other conditions, flow cytometric analysis was focused on the tri-chimera group, which revealed that virtually all (∼99%) eGFP^+^ cells were CD7^+^, confirming highly selective targeting of T cell population within PBMCs (Fig. 4C).

To evaluate T cell activation, CD25 expression was assessed in total PBMCs as a functional readout of sustained activation following AAV transduction. Conditions lacking co-stimulatory signals, including AAV6, untargeted NEO-AAV6, and NEO-AAV6 displaying single or dual targeting chimeras, showed minimal CD25 upregulation. In contrast, both CD3/CD28 bead stimulation and tri-chimera–targeted NEO-AAV6 induced a pronounced increase in CD25 expression, with the tri-chimera condition approaching, but not fully reaching, the levels observed with bead stimulation (Fig. 4D). Notably, CD25 upregulation was observed under conditions that also supported efficient gene delivery, highlighting the coupling of receptor targeting, co-stimulation, and functional transduction.

Together, these results demonstrate that programmable surface display of targeting and co-stimulatory chimeric modules enables NEO-AAV to couple T cell activation with viral delivery in a single step, improving transduction efficiency in CD7^+^ T cells within primary human PBMCs and providing a basis for downstream CAR-T cell generation.

### Generation of functional CAR-T cells via NEO-AAV delivery

Having established efficient and targeted genetic modification of CD7^+^ T cells within primary human PBMCs, we next asked whether NEO-AAV–mediated delivery could support the generation of functional chimeric antigen receptor (CAR) T cells, as a representative application of this platform. Given the negligible transduction observed with all other targeting configurations (Fig. 4), subsequent functional CAR-T studies were performed using the tri-chimera design. Primary human PBMCs from three donors were transduced with NEO-AAV6 displaying the T cell tri-chimera and encoding a CD19-directed CAR construct, as outlined in Fig. 5A, followed by assessment of CAR expression and effector function.

**Fig. 5.**
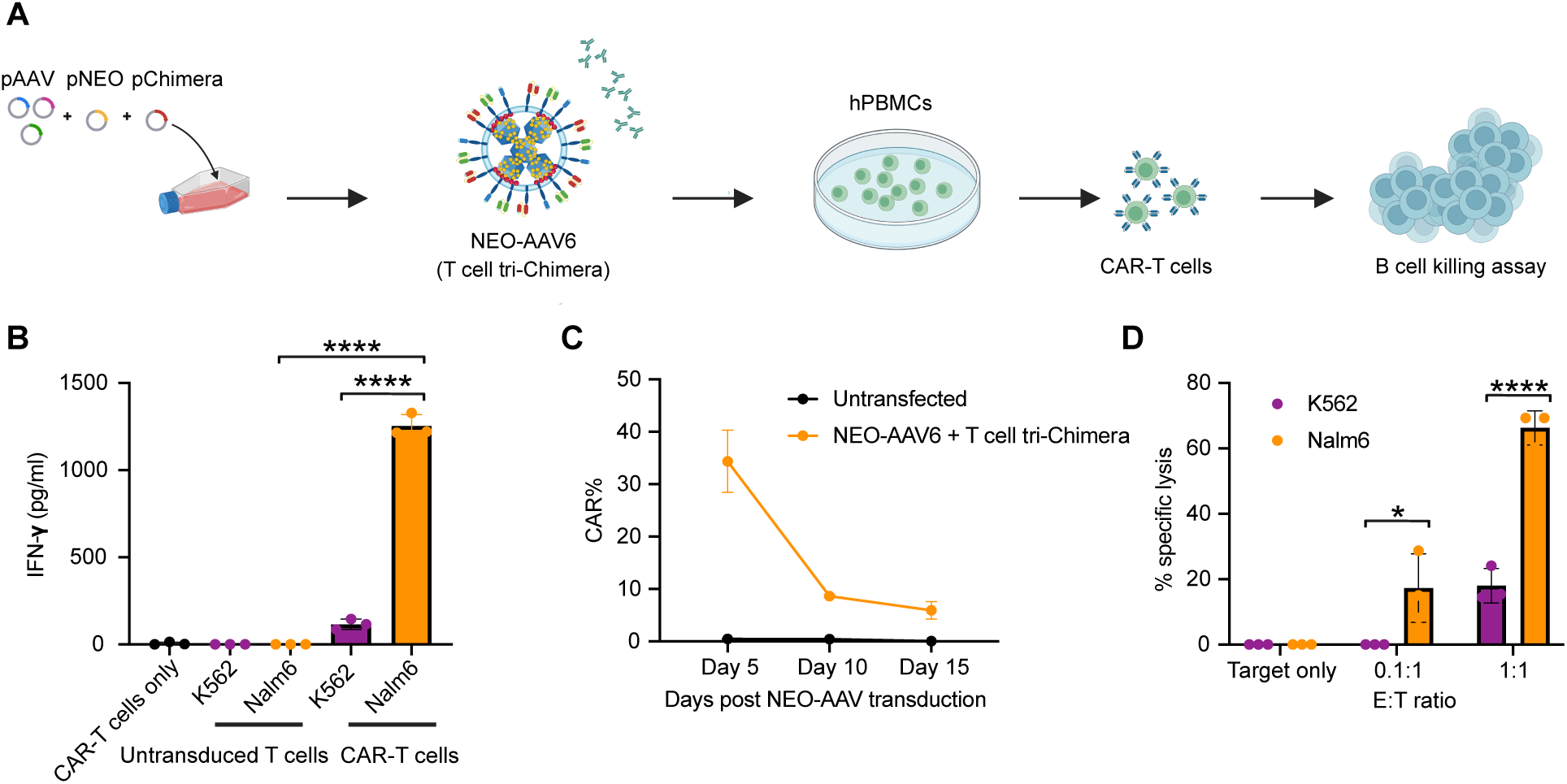
Targeted NEO-AAV enables functional CAR-T cell generation from primary human PBMCs. (**A**) Schematic workflow for CAR-T cell generation using NEO-AAV6 displaying the T cell tri-chimera and encoding a CD19-specific CAR construct. (**B**) IFN-γ secretion following co-culture of NEO-AAV–engineered CAR-T cells with CD19⁺ Nalm6 or CD19⁻ K562 target cells at an effector-to-target (E:T) ratio of 1:1. Cytokine levels were measured in the culture supernatant. Untransduced PBMCs served as controls. Data represent mean ± SD of technical duplicates from a representative donor; results were consistent across n = 3 independent donors (see Fig. 5D). Statistical significance was determined by one-way ANOVA followed by Tukey’s multiple comparisons test across treatment groups. **** p < 0.0001. (**C**) CAR expression (% CAR⁺ cells) in transduced T cells at days 5, 10, and 15 post-transduction. Data represent mean ± SD from n = 3 independent donors. (**D**) Cytotoxicity assay (% specific lysis) of CD19⁺ Nalm6 and CD19⁻ K562 target cells by NEO-AAV–generated CAR-T cells at E:T ratios of 0.1:1 and 1:1, normalized to CAR⁺ cell counts. Data represent mean ± SD from n = 3 independent donors. Statistical significance was determined using unpaired two-tailed t-tests comparing Nalm6 versus K562 within each E:T ratio. * p < 0.05; **** p < 0.0001. For all experiments, viral preparations were pre-incubated with anti-AAV neutralizing antibodies (500 ng mL⁻¹) prior to transduction at a multiplicity of infection (MOI) of 1 × 10⁶.

To evaluate antigen-dependent activation, NEO-AAV–generated CAR-T cells were co-cultured with CD19^+^ Nalm6 target cells or CD19^-^ K562 cells. CAR-T cells exhibited robust IFN-γ secretion in response to CD19^+^ Nalm6 but not CD19^-^ K562 targets, whereas untransduced PBMCs showed minimal cytokine release, indicating antigen-specific activation of NEO-AAV–generated CAR-T cells (Fig. 5B; representative donor shown).

CAR expression was assessed by flow cytometry, peaking at approximately day 5 post-transduction (∼34%) and declining to ∼6% by day 15, consistent with rapid dilution of episomal AAV genomes during the initial burst of T cell proliferation following activation ^52^ (Fig. 5C). This kinetic profile — typical of non-integrating AAV in proliferating T cells — establishes NEO-AAV + tri-chimera as a transient activation-coupled delivery module rather than a stand-alone durable engineering solution. Coupling NEO-AAV delivery with CRISPR-HDR–based CAR knock-in, in the manner of Eyquem et al. ^14^, represents a clear next step and is discussed below.

To determine whether this activation translated into cytotoxic function, we next assessed cytotoxic activity on days 6–8 post-transduction using standard effector-to-target (E:T) killing assays with E:T ratios normalized to CAR^+^ counts. NEO-AAV–derived CAR-T cells mediated efficient antigen-specific lysis of Nalm6 cells in an E:T-dependent manner (∼18% at 0.1:1 and ∼65% at 1:1), while K562 lysis remained low even at the highest E:T ratio (∼18% at 1:1) and showed no dose-dependent increase, confirming that the pronounced Nalm6 killing reflects antigen-specific cytotoxic activity rather than non-specific effector function (Fig. 5D). No measurable cytotoxicity was observed in untransduced PBMCs.

Together, these results demonstrate that NEO-AAV can generate functional CAR-T cells from primary human PBMCs through a single activation-coupled transduction step, providing an alternative to the conventional sequential bead-activation and viral-transduction workflow for ex vivo immune cell engineering. While the present study establishes proof-of-concept in an ex vivo setting, the immune-evasive and targeting capabilities of NEO-AAV suggest potential applicability to direct in vivo T cell engineering, which represents a priority direction for future development.

## Discussion

Adeno-associated virus (AAV) vectors have become a leading platform for gene delivery; however, their clinical translation remains limited by challenges including inefficient transduction of certain cell types, pre-existing immunity, and dose-dependent toxicities. Recent clinical studies have highlighted these limitations ^8, 12^, emphasizing the need for improved vector design and delivery strategies.

In this study, we present NEO-AAV as an engineered envelopment platform that re-engineers AAV biogenesis through endogenous extracellular vesicle (EV) secretion pathways, functionally converting AAV from a passively released non-enveloped virus into an EV-enveloped, actively secreted vector in HEK293FT producer cells. By coupling putative membrane targeting (PH), proposed multivalent capsid recruitment (PKD12), and ESCRT-associated release (ALG2) within a single fusion construct, NEO-AAV generates AAV-loaded vesicles that exhibit improved loading versus passive EV-AAV controls and enhanced resistance to anti-AAV neutralizing antibody.

Although naturally occurring EV-AAV particles have been reported to confer partial antibody shielding, vesicle association in these systems occurs stochastically and yields heterogeneous, poorly controlled preparations ^28–30^. Unlike passive EV-AAV systems, NEO-AAV implements a defined molecular architecture that actively redirects assembled AAV capsids into EV biogenesis pathways. This engineered envelopment process yields enhanced capsid loading, improved resistance to neutralizing antibodies under high antibody pressure, and enhanced delivery efficiency relative to conventional EV-AAV, highlighting the importance of programmed viral–vesicle coupling over passive association.

A major challenge in gene and cell therapy is efficient genetic modification of primary human T cells, which are intrinsically resistant to AAV-mediated transduction and typically require prior activation or lentiviral vectors for CAR-T manufacturing ^19, 20, 53^. Among naturally occurring serotypes, AAV6 exhibits the highest permissiveness for T cell transduction, albeit at low efficiency and high vector doses ^13–15^. By surface-displaying a CD7/CD3/CD28 tri-chimera, NEO-AAV couples activation and transduction into a single step: the tri-chimera delivers CD3/CD28 co-stimulatory signals concurrent with viral entry, activating T cells through the delivery particle itself rather than bypassing activation altogether (confirmed by CD25 upregulation in Fig. 4D). The practical advantage is workflow simplification; preservation of a naive T cell phenotype would require a fundamentally different delivery strategy and remains to be developed.

This transient expression kinetic, consistent with episomal AAV genome persistence in proliferating lymphocytes ^52^, underscores that NEO-AAV in its current form serves as an activation-coupled delivery module rather than a durable engineering solution. Stable therapeutic applications will therefore require durable transgene integration, and coupling NEO-AAV with CRISPR-mediated knock-in strategies represents a promising future direction to achieve long-term CAR expression while retaining the advantages of immune-evasive and targeted AAV delivery ^14^.

Our findings further reveal serotype-dependent differences in PKD-mediated capsid engagement. Specifically, the engineered high-affinity PKD12_EN_ variant did not improve AAV6 loading relative to PKD12_NC_ and PKD12_WT_, despite its reported enhancement of capsid binding for other serotypes ^50^. The precise PKD domain engagement of AAV6 remains incompletely characterized, and the mechanistic basis for this serotype-specific outcome warrants further investigation, potentially through structural characterization of the AAV6–PKD12 interface. Future optimization through directed evolution of serotype-specific PKD variants may therefore be required to extend this strategy across diverse AAV serotypes and therapeutic contexts. Moreover, while the present study focuses on ex vivo immune cell engineering, the ability of NEO-AAV to evade neutralizing antibodies suggests potential applicability to in vivo gene delivery scenarios, including direct immune modulation.

Together, these findings highlight the potential of NEO-AAV as a programmable platform for improving AAV delivery. By enabling controlled viral envelopment, immune evasion, and targeted delivery, this approach may offer a foundation for addressing key limitations in current gene and cell therapy strategies.

### Limitations of the current study

We wish to be explicit about what this study does and does not demonstrate. First, all neutralizing-antibody shielding data are based on a single mouse anti-AAV6 monoclonal antibody (ADK6) in HEK293FT reporter cells; shielding against pooled human IVIG and patient sera was not tested and is a priority follow-up. Second, the three-step mechanistic model (PIP2 targeting → multivalent clustering → ESCRT scission) is presented on the basis of construct design and literature precedent, but has not been formally validated by loss-of-function mutagenesis (PH PIP2-binding-dead, ALG2 EF-hand mutant, ESCRT inhibition, module truncations); such dissection is the single most important experimental extension for follow-up work. Third, CAR expression following NEO-AAV + tri-chimera delivery is transient, consistent with episomal AAV, and this work does not address durable CAR-T engineering; combination with CRISPR-HDR knock-in is identified as the natural next step. Fourth, all data are ex vivo; no in vivo efficacy, biodistribution, or immunogenicity data are presented. Fifth, T cell identity in transduced populations was primarily inferred from CD7 surface expression; in CAR-T experiments, CD2 was additionally used to identify the T cell population. However, neither CD7 nor CD2 exclusively distinguishes T cells from NK cells, and definitive lineage confirmation in future studies would require incorporation of CD56 to exclude NK cell contributions. We believe each of these gaps is tractable and outline them not as disclaimers but as a concrete roadmap for future development of this architecture.

In summary, these findings support NEO-AAV as a proof-of-concept engineered secretion route for EV-enveloped AAV and, when combined with the CD7/CD3/CD28 tri-chimera, as a single-step activation-coupled transduction module, demonstrated here in an ex vivo setting and with potential applicability to direct in vivo T cell engineering. With the mechanistic and in vivo validation outlined above, this architecture has the potential to contribute a useful building block to next-generation immune-cell engineering workflows.

## Method

### Cell culture

Human embryonic kidney 293FT (HEK293FT) (R70007, ThermoFisher) cells were maintained in Dulbecco’s Modified Eagle Medium (DMEM, 41965-039, Gibco), supplemented with 10% fetal bovine serum (FBS, 35-079-CV, Corning), 1% penicillin/streptomycin (P/S, 15-140-122, Gibco).

Primary human PBMCs were isolated from whole blood by density gradient centrifugation using Leucosep™ tubes (T2068-450EA, Greiner Bio-One) pre-filled with Ficoll-Paque PLUS (GE17-1440-03, Cytiva), according to the manufacturer’s instructions. Briefly, anticoagulated blood was diluted with PBS supplemented with 2% FBS and layered onto the separation medium, followed by centrifugation at 800–1000 × g for 10–15 min at room temperature with the centrifuge brake disengaged. The PBMC layer at the plasma–Ficoll interface was collected and washed with PBS (2% FBS) to remove residual platelets. Isolated PBMCs were maintained in X-VIVO 15 (BE02-060Q, Lonza) with 50 µM 2-mercaptoethanol (21985023, Gibco), 5% FBS and 10 mM N-acetyl l-cysteine (A9165-5G, Sigma-Aldrich).

For activation and transduction experiments, PBMCs were cultured with or without bead-based activation. Where indicated, PBMCs were stimulated with Dynabeads™ Human T-Activator CD3/CD28 (11161D, Thermo Scientific) in the presence of recombinant human IL-2 (50 U/mL, 200-02-50UG, PeproTech®), IL-7 (10 ng/mL, 200-07-10UG, PeproTech®), and IL-15 (10 ng/mL, 200-15-10UG, PeproTech®) for 2 days prior to treatment; bead-stimulated PBMCs served as a positive control for T cell activation. In tri-chimera-targeted NEO-AAV experiments, no exogenous pre-activation was applied unless otherwise stated. Following activation, whether by bead stimulation or tri-chimera-targeted NEO-AAV, cells were maintained in culture medium supplemented with IL-2 (50 U/mL) at 37 °C in a humidified atmosphere with 5% CO₂.

### Plasmids

The plasmids pcDNA3.1(+)_PH-ALG2-PKD12_WT_, pcDNA3.1(+)_PH-ALG2-PKD12_NC_, and pcDNA3.1(+)_PH-ALG2-PKD12_EN_ were synthesized by GenScript (Europe) and correspond to the pNEO_Ver 1.0_, pNEO_NC_ (EV-AAV control), and pNEO_Ver 2.0_ constructs used in this study, respectively. The plasmids pChimera_GPC3-targeting, pChimera_CD7-targeting, pChimera_CD3-targeting, and pChimera_CD28-co-stimulatory were also synthesized by GenScript; the binder sequences incorporated in these constructs are described in Section 2.9. The GPC3-targeting construct was included as a non–T cell–targeting control ^36^. The plasmid pAAV-CMV-AntiCD19-WPRE was constructed based on pAAV-CMV-GFP-WPRE. AAV packaging plasmids pAAV2/8 (Addgene plasmid #112864) and pAAV2/9n (Addgene plasmid #112865) were obtained from Addgene. All plasmids were verified by Sanger sequencing. Amino acid sequences of the NEO-AAV envelopment constructs are provided in Supplementary Data S1.

### Conventional AAV and EV-AAV/NEO-AAV production and isolation

AAV vectors were produced using either a triple-plasmid system (AAV8 and AAV9; vector:rep/cap:helper = 1:1:1) or a dual-plasmid system (AAV6; transfer:packaging = 1:2), with a total of 30 μg DNA per T175 flask. HEK293FT cells were cultured to ∼80% confluency and transfected using Lipofectamine® 3000 Transfection Reagent (L3000008, Invitrogen) at a ratio of 1 μg DNA to 2 μL reagent according to the manufacturer’s instructions. Following transfection, cells were maintained for 72 h before virus harvest.

For virus collection, both cells and culture supernatant were harvested. Cells were pelleted and subjected to three freeze–thaw cycles to release intracellular AAV particles, followed by Benzonase (50 U/mL) treatment and detergent-mediated lysis. The clarified cell lysate was combined with the culture supernatant.

AAV particles were precipitated using polyethylene glycol (PEG8000) in the presence of NaCl and further purified by aqueous two-phase partitioning with PEG8000 and ammonium sulfate. After phase separation, the virus-containing aqueous fraction was collected, dialyzed against PBS to remove salts, and concentrated using centrifugal filter units. The final preparation was sterile-filtered (0.22 μm) and stored at −80 °C until use.

For EV-AAV and NEO-AAV production, 10 μg of pNEO construct (pNEO_Ver 1.0_, pNEO_NC_ or pNEO_Ver 2.0_, depending on the experimental condition) was added to the transfection mixture, bringing the total DNA to 40 μg per T175 flask. For T cell-targeted NEO-AAV production, additional targeting plasmids (24 μg total) were included, bringing the total DNA to 64 μg per T175 flask, either as single constructs or in multiplexed combinations at equal ratios (1:1 or 1:1:1). At 72 h post-transfection, the culture supernatant was collected and subjected to sequential centrifugation steps for removal of cells and debris. Briefly, the supernatant was centrifuged three times at 500 × g for 5 min at 4 °C to remove cell debris, followed by centrifugation at 2,000 × g for 30 min at 4 °C to remove smaller debris and apoptotic bodies. The clarified supernatant was then filtered through a 0.45 μm filter to eliminate residual cellular contaminants. Extracellular vesicles containing AAV (EV-AAV/NEO-AAV) were pelleted by ultracentrifugation at 135,000 × g for 90 min at 4 °C. As ultracentrifugation may co-pellet free AAV particles, the EV pellet was resuspended in PBS and further purified using qEVsingle 70nm size exclusion chromatography columns (SP2, Izon Science) following the manufacturer’s instructions. AAV genome titers were determined by quantitative PCR (qPCR) as previously described ^37^. Briefly, to ensure accurate titration of EV-AAV containing protein and lipids, AAV genomes were purified using the High Pure Viral Nucleic Acid Kit (11858874001, Roche) prior to qPCR. Residual plasmid DNA from transfection was first removed by treating 5 μL of EV-AAV sample with DNase I (1 μL DNase I, 5 μL 10× buffer, 39 μL water) at 37°C for 1 h, followed by inactivation at 75°C for 15 min. Genome titers were then determined using a TaqMan qPCR assay targeting the polyA region of the transgene cassette.

### Cryo-electron microscopy (Cryo-EM)

Cryo-EM analysis was performed by ATEM Structural Discovery (Germany). Briefly, 4 µL of sample was applied to holey carbon grids and incubated under controlled temperature (4 °C) and humidity conditions using a Vitrobot Mark IV (Thermo Fisher Scientific). Grids were blotted and rapidly vitrified by plunge-freezing in liquid ethane and stored in liquid nitrogen prior to imaging.

Cryo-EM imaging was carried out on a 200 kV Thermo Fisher Scientific Glacios transmission electron microscope equipped with a Falcon IV direct electron detector. Images were acquired under low-dose conditions at approximately 45,000× magnification using dose fractionation. Raw movies were motion-corrected and frame-aligned prior to downstream analysis.

### Nanoparticle Tracking Analysis

The size and particle concentration of EVs were assessed with nanoparticle tracking analysis (NTA) (NS500, Malvern Nanosight). EVs were suspended in PBS and measured in triplicate with individual measurements of 30 seconds at camera level 15. Analysis was performed with NTA software 3.3 with a minimal track length of 10, detection threshold 5, and screen gain 1. The cut-off values for reliable measurement were between 20.0 and 100.0 particles/frame ^38^.

### Western blotting

Cell lysates were prepared using cOmplete™ Lysis-M EDTA-free (4719964001, Roche) according to the manufacturer’s instructions. EV-AAV/NEO-AAV samples were lysed in RIPA lysis buffer (1:10, 20-188, Sigma) supplemented with Protease/ Phosphatase inhibitor cocktail (1:100, 5872S, Cell Signaling Technology) and stored on ice for 30 min. Next, EV-AAV/NEO-AAV samples were additionally centrifuged at 14,000 × g for 10 min at 4 °C to remove residual membrane debris, and the supernatant was collected. The lysed EVs and cell lysates were stored at -80 °C.

Protein concentrations were determined via the Micro BCA Protein Assay Kit (23235, ThermoFisher). Samples were mixed with NuPAGE™ Sample Reducing Agent (50 mM, NP0004, Invitrogen Corp) and NuPAGE LDS Sample Buffer (NP0007, Invitrogen), incubated at 70 °C for 10 min. Samples were separated on Bolt™ 4–12% Bis-Tris Plus Gel (NW04125BOX, ThermoFisher Scientific), with a PageRuler Plus Prestained Protein Ladder (26619, ThermoFisher Scientific) at 200 V for 35 min and transferred to PVDF membranes (IPVH00010, Merck). The membranes were blocked for 1 h in Intercept® (TBS) Blocking Buffer (927-60003, LICORbio).

Primary antibodies included mouse anti-AAV VP1/VP2/VP3 (1:5000 dilution, Progen, B1, AFDye™ 488 Conjugate), rabbit anti-TSG101 (1:1000 dilution, ab30871, Abcam), mouse anti-flag (1:1000, F1804-200, Sigma-Aldrich), and mouse anti-β-actin (1:5000, Cell Signaling Technology, clone 8H10D10). Secondary antibodies included Goat anti-mouse antibody (1:1000 dilution, P0447, dako) and Goat anti-rabbit antibody (1:2000 dilution, P0448, dako).

### Nano-ffow cytometry analysis

Extracellular vesicles (EVs) were characterized using Nano-flow cytometry (NanoFCM, NanoAnalyzer, NanoFCM Inc.) to assess particle size, concentration, and AAV cargo loading.

For size distribution and particle concentration measurements, EV samples were diluted to approximately 2 × 10⁸ particles mL⁻¹ in 10 mM HEPES buffer and analyzed directly using the NanoAnalyzer.

For internal AAV cargo detection, EVs were first fixed by mixing with an equal volume of 8% paraformaldehyde and incubating for 30 min at room temperature. After removal of fixative using SEC columns, EVs were permeabilized with an equal volume of 2× NanoFCM permeabilization buffer and incubated on ice for 30 min. Permeabilized EVs were then incubated with CaptureSelect™ Alexa Fluor™ 647 Anti-AAVX conjugate (7233522100, Thermo Fisher) at a ratio of 10 μL antibody per 90 μL EV suspension for 30 min at 37 °C. Excess antibody was removed by size exclusion chromatography (qEVsingle, Izon Science), and samples were diluted to ∼2 × 10⁸ particles mL⁻¹ prior to NanoFCM analysis.

Fluorescence signals were used to determine the proportion of EVs positive for AAV cargo. Auto-fluorescence signals were evaluated in unstained controls and used to guide channel selection and labeling strategies.

### Neutralization assay

HEK293FT cells were seeded at a density of 1 × 10⁴ cells per well in 96-well plates (655075, Greiner CELLSTAR) and cultured overnight under standard conditions. EV-AAV/NEO-AAV preparations were incubated with increasing concentrations of anti-AAV6 monoclonal neutralizing antibody (ADK6, 0, 5, 50, and 500 ng mL⁻¹; Progen) at 37 °C for 1 h. Following neutralization, mixtures were added to pre-seeded HEK293FT cells at an MOI of 1.5 × 10⁴ and incubated for 72 h. Transduction efficiency was assessed by eGFP fluorescence imaging (EVOS M5000, Invitrogen) and quantified by flow cytometry (CytoFLEX, Beckman Coulter).

### T cell-targeting binders

CD7-specific nanobodies were generated through immunization and phage display selection. Briefly, two llamas were immunized with lipid nanoparticle (LNP)-formulated mRNA encoding full-length human CD7. Peripheral blood lymphocytes were isolated, and VHH-encoding sequences were amplified and cloned into a phagemid vector to construct a phage display library. For binder selection, live-cell biopanning was performed using CD7^+^ target cells, with CD8^+^ cells used for negative selection. Phage libraries were iteratively enriched through three rounds of selection. Individual clones were screened for CD7 binding by flow cytometry using target cells and primary human PBMCs. Positive clones were sequenced, and selected VHHs were expressed in E. coli and purified via His-tag affinity chromatography. Binding affinity of the lead VHH used in this study is summarized in Supplementary Fig. S1.

CD3-specific scFvs (OKT3 parental clone) were obtained through a commercial antibody engineering service (Creative Biolabs), including sequence synthesis, recombinant expression, and affinity purification. The CD28 co-stimulatory binder sequence was derived from Hamilton et al. ^19^ and incorporated into the chimera construct during plasmid synthesis (GenScript).

### Flow cytometry analysis

For CAR expression analysis, CD19 CAR was detected using CD19 CAR detection reagent (Miltenyi Biotec, REA675, APC, 1:400), and T cells were identified using anti-hCD2-BV605 (clone S5.2, BD Pharmingen, 745205, 1:1000). T cell activation status was assessed using anti-hCD25-PE-CF594 (BD Pharmingen, 562403, 1:200). For T cell specificity experiments using PBMCs, a ZombieNIR (BioLegend) viability dye and Fc Block (BD, 564220, 1:200) were applied prior to staining. T cells were identified using anti-hCD7-APC (BD Pharmingen, 561604, 1:500).

### Microscopy analysis

Microscopy analysis was performed using the EVOS FL Cell Imaging System (Life Technologies). Live cell brightfield and fluorescence imaging was used to visualize eGFP expression and assess cell morphology following treatment. Images were acquired in brightfield and GFP channels using identical acquisition settings within each experiment.

### Functional T cell analysis

A luciferase-based killing assay was performed to evaluate the cytotoxic function of CAR T cells generated via tri-chimera-targeted NEO-AAV delivery. CAR expression was confirmed prior to the assay, and effector-to-target (E:T) ratios were normalized to CAR^+^ cell counts. On days 6–8 post-transduction, CAR T cells were co-cultured with 3,000 target cells (Nalm6, CD19^+^; K562, CD19^−^) at varying E:T ratios (0:1, 0.1:1, and 1:1) for 24 h. Luciferin (E501, Promega) was then added to each well and luminescence was measured using a SpectraMax iD3 microplate reader. Untreated PBMCs from the same donor were used as negative controls.

### Interferon-gamma Elisa

Interferon-gamma (IFN-γ) levels were quantified in cell culture supernatants using the Human IFN-γ Quantikine ELISA Kit (DIF50C, RCD Systems) according to the manufacturer’s instructions. Cell culture supernatants were collected 3 days after co-culture with target cells, cleared by centrifugation at 500 × g for 5 min, and diluted 20-fold in Calibrator Diluent RD5P prior to analysis. Absorbance was measured at 450 nm with correction at 570 nm using a Multiskan FC microplate reader (Thermo Scientific). IFN-γ concentrations were calculated from a standard curve ranging from 15.6 to 1000 pg mL⁻¹ using a four-parameter logistic curve fit.

### Statistical test

Statistical analyses were performed using GraphPad Prism (v.9.3). Data are presented as mean ± s.d. or mean ± SEM, as indicated in the figure legends. For experiments involving biological replicates (e.g., independent PBMC donors or EV-AAV preparations), variability is generally reported as SD. Comparisons between two groups were performed using a two-tailed unpaired t-test (with Welch’s correction where appropriate), and comparisons among multiple groups were analyzed using one-way ANOVA followed by Tukey’s or Dunnett’s multiple comparisons test, as specified. Exact P values and statistical tests are indicated in the corresponding figure legends. p < 0.05 was considered statistically significant.

## Supporting information

CD7 VHH Validation

## Acknowledgements

The authors thank Mischa Klerk for his experimental support. The authors gratefully acknowledge ATEM Structural Discovery (Germany) for their support with cryo-electron microscopy (cryo-EM) analysis. We also thank NanoFCM for assistance with Nano-flow cytometry measurements. Their technical expertise and support were essential for the successful completion of this study. Schematic illustrations have been created with BioRender.com. During the preparation of this manuscript, the authors used Claude to assist with language editing and proofreading; all content was subsequently reviewed and edited by the authors, who take full responsibility for the published work.

Author Contributions

S.X. and Q.Y. contributed equally to this work. S.X. conceived the NEO-AAV platform, designed and performed experiments including plasmid construct generation, vector production and characterization, reporter assays, and functional delivery studies, and wrote the manuscript. Q.Y. contributed to experimental design, performed experiments, and co-wrote the manuscript. K.Q. assisted with manuscript preparation and figure illustration. C.S.B. provided technical support. L.V.J., B.Y., G.Y., S.C.A.d.J., J.X., P.V., G.J.J.B., R.M.S., and P.D. provided scientific input and critically reviewed the manuscript. J.P.G.S. and Z.L. supervised the project, secured funding, and critically reviewed and edited the manuscript. All authors read and approved the final manuscript.

## Data Availability

The datasets used and/or analyzed in the current study are available from the corresponding authors upon reasonable request.

## Competing interests

S.X., J.P.G.S., and Z.L. have filed a patent related to the NEO-AAV platform described in this work. R.M.S. and Z.L. are employees of NanoCell Therapeutics, a company developing lipid nanoparticles for in vivo CAR-T cell therapy. G.J.J.B. reports ownership interest in PacingCure B.V. The remaining authors declare no competing interests.

## Funding

J.P.G.S. is supported by ZonMw Psider-Heart (10250022110004), NWO-TTP HARVEY (2021/TTW/01038252), H2020- TOP-EVICARE (#101138069) and VIA-EVICARE (#101212624) of the European Research Council (ERC), Health-Holland 2022TKI2306 (EV-PROTECT), Netherlands Heart foundation 01-003-2024-0455 (EVOLVE), and ERA for Health Cardinnov (RESCUE- 2024/KIC/01627794).

## Notes

### Competing Interest Statement

The authors have declared no competing interest.

### Summary of Updates

This version of the manuscript has been revised to update the following: Manuscript text revised for clarity and language throughout. Results section updated to clarify transduction efficiency reporting in Figure 4. Methods section expanded with additional experimental details. Author list and affiliations updated. Competing interests statement updated to reflect patent filing.

