## Supplementary material for "NEO-AAV: an engineered extracellular vesicle-enveloped AAV platform for activation-coupled T cell transduction and CAR-T cell generation": CD7 VHH Validation

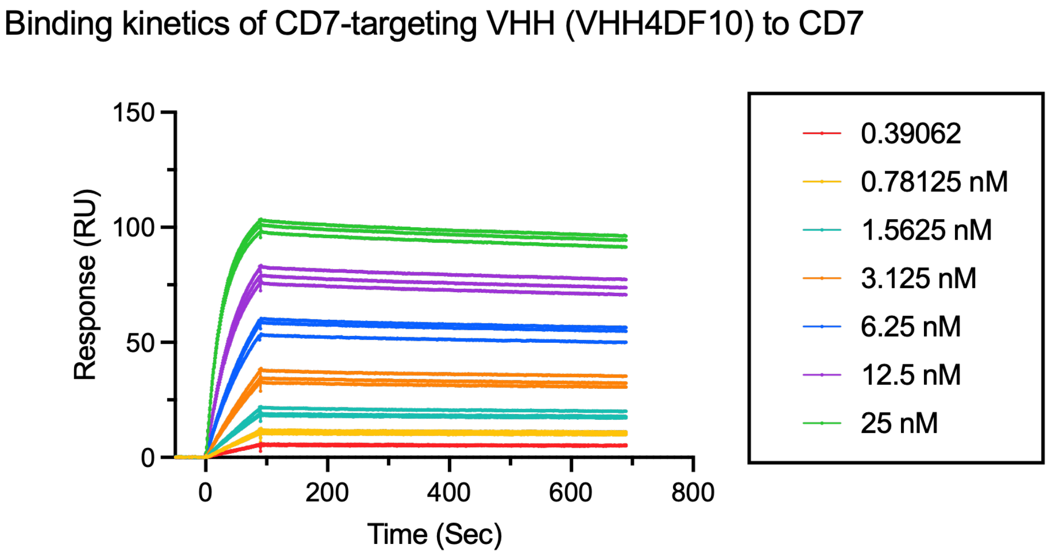


**Fig. S1 | Binding kinetics of CD7-targeting VHH (VHH4DF10) to CD7.**

Representative sensorgrams showing concentration-dependent binding of VHH4DF10 to CD7, measured by surface plasmon resonance (SPR). Serial dilutions of VHH4DF10 (0.39–25 nM) were injected, and real-time association and dissociation responses were recorded. Kinetic parameters were determined by global fitting using a 1:1 binding model, yielding an association rate constant of $1.42 \times 10^{6}\ M^{-1}\ s^{-1}$, a dissociation rate constant of $1.07 \times 10^{-4}\ s^{-1}$, and an equilibrium dissociation constant of 0.075 nM.
